# micrIO: An Open-Source Autosampler and Fraction Collector for Automated Microfluidic Input-Output

**DOI:** 10.1101/655324

**Authors:** Scott A. Longwell, Polly M. Fordyce

## Abstract

Microfluidic devices are an empowering technology for many labs, enabling a wide range of applications spanning high-throughput encapsulation, molecular separations, and long-term cell culture. In many cases, however, their utility is limited by a ‘world-to-chip’ barrier that makes it difficult to serially interface samples with these devices. As a result, many researchers are forced to rely on low-throughput, manual approaches for managing device input and output (IO) of samples, reagents, and effluent. Here, we present a hardware-software platform for automated microfluidic IO (micrIO). The platform, which is uniquely compatible with positive-pressure microfluidics, comprises an ‘AutoSipper’ for input and a Fraction Collector for output. To facilitate wide-spread adoption, both are open-source builds constructed from components that are readily purchased online or fabricated from included design files. The software control library, written in Python, allows the platform to be integrated with existing experimental setups and to coordinate IO with other functions such as valve actuation and assay imaging. We demonstrate these capabilities by coupling both the AutoSipper and Fraction Collector to a microfluidic device that produces beads with distinct spectral codes, and an analysis of the collected bead fractions establishes the ability of the platform to draw from and output to specific wells of multiwell plates with no detectable cross-contamination between samples.

## 1 Introduction

Microfluidic devices are powerful tools for biology, chemistry, and medicine, with applications including biomolecular characterization, ^1,2^ cell encapsulation and culture, ^3–5^ particle synthesis, ^6,7^ and diagnostics. ^8^ In theory, their miniature scale allows a researcher to integrate processes that span several lab benches into a single device with minimal cost and sample requirements. In reality, the extensive infrastructure required to convey reagents, samples, and analytes into and out of devices often renders a ‘lab-on-a-chip’ more of a ‘chip-in-a-lab.’ ^9–11^ A flexible interface allowing samples in common labware formats to be introduced into and collected from microfluidic devices would help surmount the ‘world-to-chip’ barrier and make it possible for more laboratories to benefit from the full potential of on-chip automation. ^12–15^

Treating a microfluidic device as a processing module, this challenge can be described as microfluidic input-output (IO). Low-throughput, serial IO is easily handled through manual exchange of vessels connected to a device. However, long-term operation of a device with many inputs and outputs (multiplexed IO) quickly becomes tedious and carries an increased risk of user error. While integrated on-chip valves aid in serial multiplexing and demultiplexing by enabling automated selection of inputs and outputs,14,16–19 they do not scale well beyond a dozen inputs and outputs, as each additional valve requires researchers to connect additional control and flow lines during setup. An alternate approach that moves the IO interface off-chip could allow samples to be introduced from standardized multiwell plate formats (e.g. 96-well plates) and make high-throughput multiplexed IO trivial to implement. Autosamplers (for input) and fraction collectors (for output) are well-established tools for interfacing plates with a variety of analytical chemistry instruments in an automated fashion. However, their adoption in microfluidic applications has been limited, largely because commercial solutions designed for microfluidic applications are expensive and their closed-source control software makes programmatic integration into existing workflows difficult. Moreover, in contrast to the volumetric sample flows used in applications like high-performance liquid chromatography (HPLC), microfluidic devices that incorporate on-chip valves are frequently run with positive, pressure-driven flow to avoid over-pressuring and delaminating the device. The use of pressure-driven flow also leverages existing infrastructure required to control integrated pneumatic valves to drive fluid flow through the device.^16,20^

Here, we describe a low-cost, open-source platform for high-throughput microfluidic IO (micrIO). It comprises an ‘AutoSipper’ that allows for high-throughput introduction of samples from a multiwell plate into a microfluidic device via pressure-driven flow, a Fraction Collector for high-throughput sample collection from a microfluidic device into a multi-well plate or vial rack, and an open-source Python control-software suite. All hardware components are readily available as either used parts on eBay, from suppliers like McMaster-Carr or Amazon, or included as design files for 3D-printing or laser-cutting.^†^ Our control software, written in Python, is available as the pip-installable package *acqpack*, with source code available as a public repository on GitHub. The hardware and software are both modular, allowing end users to integrate additional components or adapt micrIO to better fit their needs. To validate the platform, we connected the AutoSipper and Fraction Collector to a microfluidic droplet/bead generator capable of producing spectrally encoded polymeric beads from aliquoted LN-prepolymer mixtures. ^7,21^ This experiment demonstrated the ability of the platform to a) introduce a sequence of 9 coded LN-prepolymer mixtures from a 96-well plate into a microfluidic bead generator and b) collect the coded bead batches in separate output vials with no detectable crosstalk. We anticipate that this platform will prove broadly useful to researchers who routinely employ either simple or valved microfluidic devices and are currently bottle-necked by serial sample introduction and collection.

## 2 Materials and methods

### 2.1 Assembly of AutoSipper and Fraction Collector

In the below assembly description, all custom 3D-printed or laser-cut components were designed in Autodesk Fusion360. 3D-printed components were printed from ABS filament on a Stratasys uPrint Plus. Laser-cut components were cut from nominally 1/4” acrylic sheets on a Universal Laser Systems laser cutter. Detailed information for constructing or modifying the platform, such as a complete parts list and full CAD model, is contained within the mircIO GitHub repository.^†^

To assemble the micrIO platform, the structural frame was first constructed from 80/20 T-slot framing and brackets (McMaster-Carr) and mounted on a 6” × 6” optical bread-board (Thorlabs MB6). To form the base of the AutoSipper and Fraction Collector, two XY-stages (Applied Scientific Instrumentation OEM MS-2000), salvaged from a decommissioned Illumina GAIIx and purchased on eBay, were mounted on 80/20 arms of the structural frame with 80/20 brackets. Each stage was then affixed with 3D-printed holders to accept standard ANSI/SLAS plates. The AutoSipper stage additionally received a laser-cut brace plate (to help multiwell plates resist deflection during sampling), as well as a 3D-printed vial holder for the placement of up to 4 scintillation vials.

The AutoSipper Z-assembly was attached to the vertical rail of the structural frame. It consists of a laser-cut backplate to which several components were mounted. The rotary motion of a stepper motor with optical homing sensor (Lin Engineering CO-4118S-09; also salvaged from a GAIIx) was adapted to drive a carriage up and down a linear way via a leadscrew, anti-backlash nut, and crossbar. A 3D-printed sampler arm was attached to the carriage to enable vertical movement of its end effector, a dual-lumen sampling needle. The dual-lumen needle was fabricated by pushing two 22G sample needles (trimmed to length) through a Luer end cap, reinforcing the punctured area with epoxy, and affixing a grommet (for 3/8” hole) to the end of the cap with epoxy.

The Fraction Collector dropper assembly consists of a 3D-printed arm and tubing holder/sheath nozzle that slides into position along the arm. The arm was attached directly to the vertical rail of the structural frame. A laser-cut tube holder for lashing a pressurized sheath fluid vessel (15 mL Falcon tube) with elastic (*e.g.* Tygon tubing) was also affixed directly to the structural frame.

To control the platform, the XY-stages were connected to separate ASI LX-4000 stage controllers (also salvaged from a GAIIx). The stepper motor has an integrated controller driver and does not require an additional controller. Each controller was connected to a PC via USB-serial adapters.

### 2.2 Fabrication of microfluidic bead generator

Molding masters and PDMS devices for the valved microfluidic T-junction bead generator design used here were fabricated according to a previously described protocol. ^22^ Briefly, master mold wafers for casting the control and flow layers of the PDMS devices were prepared by multilayer photolithography using AZ50 XT (Capitol Scientific) and SU-8 (Microchem) photoresists according to the manufacturer’s specification. Two-layer devices with integrated pneumatic valves were then fabricated from these molding masters by casting PDMS (R.S. Hughes RTV615).

### 2.3 Preparation of LN-prepolymer mixtures

Prepolymer mixtures containing lanthanide nanoparticles were prepared as in Nguyen, *et. al.* ^21^ using lanthanide yttrium orthovanadate nanophosphors (LNs) synthesized and wrapped in polyacrylic acid as described previously. ^7^ Briefly, 1.842 mL of a prepolymer master mix was prepared by combining 942 µL polyethylene glycol diacrylate 700 (PEG-DA; Sigma-Aldrich 455008), 724 µL Milli-Q water, 110 µL 100 mM HEPES (pH 6.8), and 66 µL of a solution containing 39.2 mg/mL of the photoinitiator lithium phenyl-2,4,6-trimethylbenzoylphosphinate (LAP; Sigma-Aldrich 900889) in 100 mM HEPES. To create 400 µL each of four LN-prepolymer mixtures (Eu, Dy, Sm, and blank), 335 µL of the master mix was combined with 65 µL the appropriate LN suspension (either 50 mg/mL Eu:YVO_4_, 50 mg/mL Dy:YVO_4_, 50 mg/mL Sm:YVO_4_, or an equal volume of Milli-Q water). After mixing by pipette, each LN-prepolymer mixture was spun through a 0.45 µm PVDF filter (Millipore UFC40HV) to remove particulates and then dispensed into a 96-well skirted plate (Fisherbrand 14230238) as follows to a volume of 125 µL per well: Eu to A01, D06, H10; Dy to A02, D07, H11; Sm to A03, D08, H12; and blank to B01, B02, B03. The plate was sealed with an adhesive aluminum foil seal (ThermoFisher AB0626).

### 2.4 Bead synthesis device setup

All valve control ports were connected via a blunt steel pin (O.D. 0.025 in, I.D. 0.013 in; New England Small Tube NE-1310-02) and Tygon ND-100-80 tubing (O.D. 0.06 in, I.D. 0.02 in; Fisher Scientific 14-171-284) to a control manifold, ^20^ then primed with water to dead-end fill control lines. The oil flow inlet was similarly connected via a steel pin and Tygon tubing to a pressurized vessel containing 2% w/w ionic krytox (Miller Stephenson 157 FSH) in HFE-7500 (3M Novec 7500); the wash inlet was connected via a steel pin and Tygon tubing to a pressurized vessel containing 50% v/v ethanol for device flushing. The oil vessel and wash vessel were each pressurized with a microfluidic control system (Fluigent MFCS-EZ) for computer-scriptable pressure adjustment. The end of a 3 mm liquid light guide connected to a UV spot curing system (Dymax 41015) was positioned above the outlet channel (5 mm from the surface of the PDMS) for polymerization of droplets into solid beads.

### 2.5 AutoSipper and Fraction Collector setup and operation

To prepare the AutoSipper, a 96-well plate (Fisherbrand 14230238) containing LN-prepolymer mixtures was placed on the deck. In addition, three 20-mL scintillation vials (Wheaton 986731) were set in the deck’s vial holder: an empty waste vial, a strong wash vial containing 20 mL of isopropyl alcohol, and a weak wash vial containing 20 mL of water. The headspace port of the dual-lumen needle was connected to an MFCS-EZ channel with Tygon tubing. The sample port of the dual-lumen needle was connected via 50 cm of of PEEK tubing (O.D. 510 µm, I.D. 255 µm; Zeus Industrial Products, custom order) to one of two bead generator inlets.

To prepare the Fraction Collector, a machined 48-vial rack holding 5 mL fritted peptide synthesis vessels (Torviq SF-0500) with 500 µL dimethylformamide (DMF; Sigma-Aldrich) in each was placed on the deck’s plate slot. The outlet sheath nozzle was connected via a steel pin and Tygon tubing to a vessel with the headspace pressurized by an MFCS-EZ channel. The outlet of the bead generator was connected via 40 cm of PEEK tubing to the Fraction Collector’s dropper assembly. All details regarding device operation, including scripting routines used, are available as a Jupyter notebook in the micrIO GitHub repository.^†^

### 2.6 Bead imaging and analysis

Each collected fraction of beads was washed sequentially with 3 × 5 mL of DMF, 3 × 5 mL of ethanol, and 3 × 5 mL of phosphate-buffered saline with 0.1% v/v Tween-20 (PBS-T) before being resuspended in 200 µL of PBS-T. Aliquots (∼ 20 µL) of beads were placed on a glass slide, covered with a quartz coverslip, and imaged on a modified Nikon Ti-E microscope with UV-254 nm excitation and 9 bandpass emission filters as described previously. ^21,23,24^

Bead images, each consisting of 9 lanthanide channels, were analyzed with a Python analysis pipeline (included as a Jupyter notebook within the micrIO GitHub repository^†^) that used processing functionality from *skimage, cv2*, and *scipy*. Briefly, images were loaded into memory as *numpy* arrays using *tifffile*. To process a single image, pixel intensities were summed across the 3 channels that best distinguished the 3 LNs (572 nm, 620 nm, 650 nm) to produce an image that was then (a) Otsu thresholded to separate background from fore-ground regions and (b) morphologically opened and eroded to remove bright pixels, dust, and edge artifacts. Foreground regions of this summed, thresholded image were then analyzed by a peak finding algorithm to identify bead centers, which were in turn used to a seed a watershed segmentation. The watershed segmentation assigned all pixels to regions corresponding to putative bead regions (or background), allowing calculation of bead region properties such as pixel area and per channel median intensity. This process was applied to all acquired images to yield a complete list of bead regions.

To correct for positional dependence in observed median bead intensity *I*_*obs*_(*x, y*) within each channel, the parameters of a parabolic flat-field equation *S*(*x, y*) were estimated by fitting to the median intensities of beads whose putative LN was brightest in that channel:

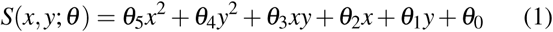

For every bead, the corrected median intensity *I*_*corr*_ in each channel was then estimated as:

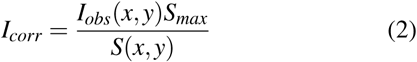

To produce the plots in Fig. 5D and E, we limited analysis to bead regions within 368 px of the image center as this encircled area corresponded to the disk of illumination from the UV light source.

## 3 Results and discussion

A microfluidic multiplexed IO platform with broad utility should be accessible, flexible, reliable, and useful for a wide variety of tasks. To meet these requirements, we developed a modular open-source microfluidic IO platform composed of an ‘AutoSipper’, which allows for serial introduction of samples from multiwell plates into microfluidic device inputs, and a ‘Fraction Collector’, which allows for serial collection from microfluidic device outputs to another multiwell plate or tube rack (Fig. 1A). The AutoSipper was designed to be compatible with microfluidic systems in which samples are introduced via pressure-driven flow and therefore assumes simple pressure control components are available (*e.g.* pressure-regulated house air, a microfluidic flow controller, or a pneumatic control manifold). All modules are comprised of low-cost hardware components that are either widely available or easily fabricated and the overall assembly can be adapted as necessary for a variety of tasks. To facilitate widespread adoption, the supplemental GitHub repository^†^ includes all information necessary to assemble and control micrIO, including a detailed CAD rendering, design files for 3D-printed or laser cut components, a calibration guide, and software documentation. For detailed information on how to build a microfluidic pneumatic control manifold, please see our previously published manuscript ^20^ as well as a low-cost Arduino-based controller for interfacing the manifold with a PC. ^25^

**Fig. 1.**
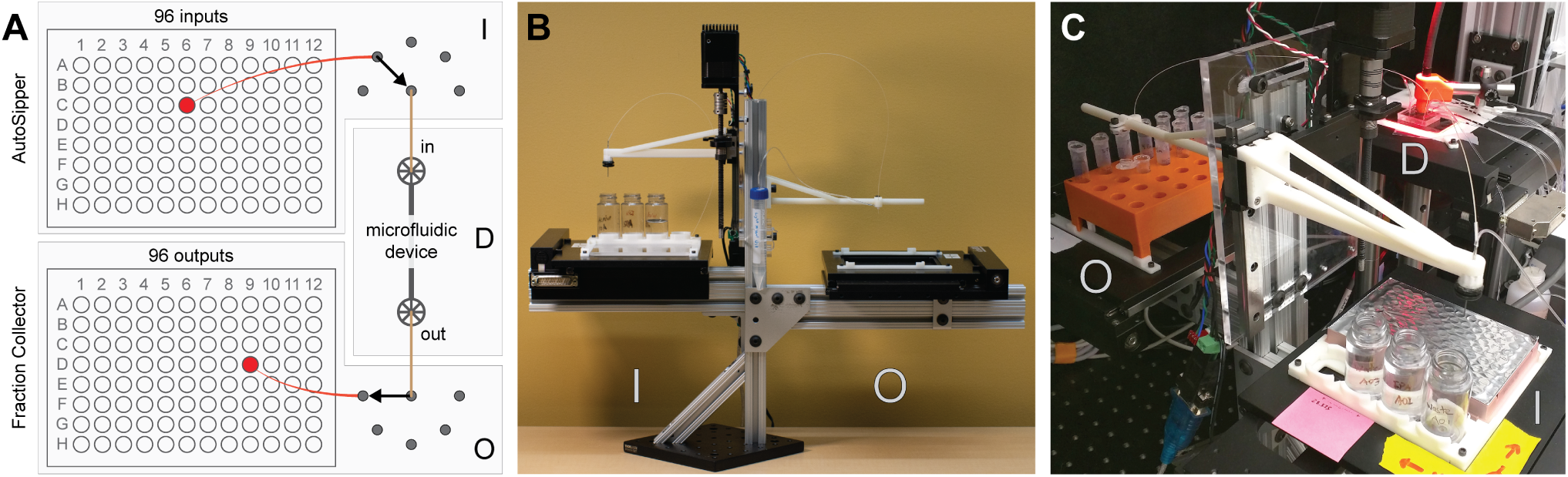
Overview of the micrIO platform. **(A)** Cartoon schematic depicting a general experimental setup in which the ‘AutoSipper’ sampler (I) introduces an input sample from a multiwell plate into a microfluidic device (D) while the Fraction Collector (O) outputs effluent from the microfluidic device into a second multiwell plate. **(B)** Photograph showing overall platform with AutoSipper (I) and Fraction Collector (O) labeled. **(C)** Photograph depicting AutoSipper (I) and Fraction Collector (O) connected to an experimental setup in which a valved microfluidic device (D) is monitored on a microscope.

### 3.1 Structural frame

To eliminate long stretches of tubing that increase dead volumes and wash times, the AutoSipper and Fraction Collector are mounted on opposite sides of a standalone 80/20 frame that can also be used for flexible mounting of a microfluidic device if desired (Fig. 1B). This frame can either be affixed directly to an optics table (Fig. 1C) or to a small optics breadboard for free movement on a benchtop (Fig. 1B). The balanced, cantilevered arms that support the AutoSipper and Fraction Collector stages are sturdy yet avoid increasing the effective platform footprint, and the single vertical rail allows height adjustment to ensure compatibility with existing experimental platforms, such as microscopes (Fig. 1C). The use of 80/20 further simplifies mounting of additional vessels or alternate equipment as needed.

### 3.2 AutoSipper

High-throughput multiplexed sample input into microfluidic devices requires the ability to: (1) serially move to sample locations, (2) interface with sample wells in a repeatable manner, and (3) push sampled fluid into a microfluidic device for downstream experiments and processing. For applications where contamination between samples must be minimized, the AutoSipper must also allow complete washing of sample lines (and potentially device channels) between each injection. Finally, truly high-throughput multiplexed sample input requires that the process be fully automated, requiring no user intervention after initial programming. To meet these objectives, we designed the AutoSipper with a gasketed dual-lumen needle attached to a 3D-printed arm mounted on a computer-controlled Z-assembly (Fig. 2A, Fig. 3). For serial addressing of samples distributed throughout a multiwell plate, the entire sample plate is mounted on an computer-controlled XY-stage.

**Fig. 2.**
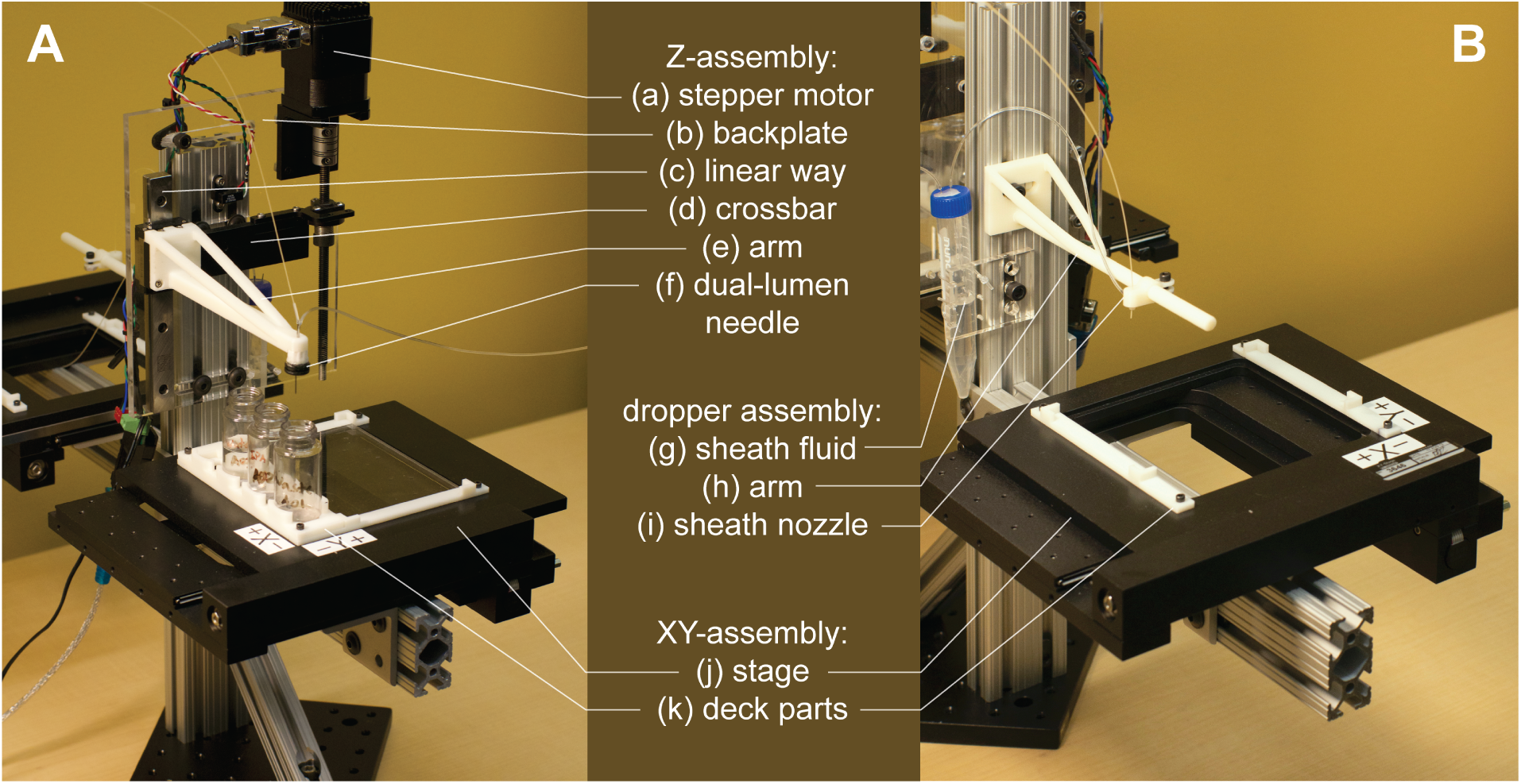
Labeled images of the AutoSipper and Fraction Collector. **(A)** The AutoSipper comprises an XY-stage with 3D-printed and laser-cut deck components for affixing multiwell plates and wash vials, as well as a Z-assembly for sampling from wells with the dual-lumen needle. **(B)** The Fraction Collector includes a similar XY-assembly, as well as a dropper assembly which enables optional rinsing of the collection tubing outlet with a sheath fluid (such as water).

**Fig. 3.**
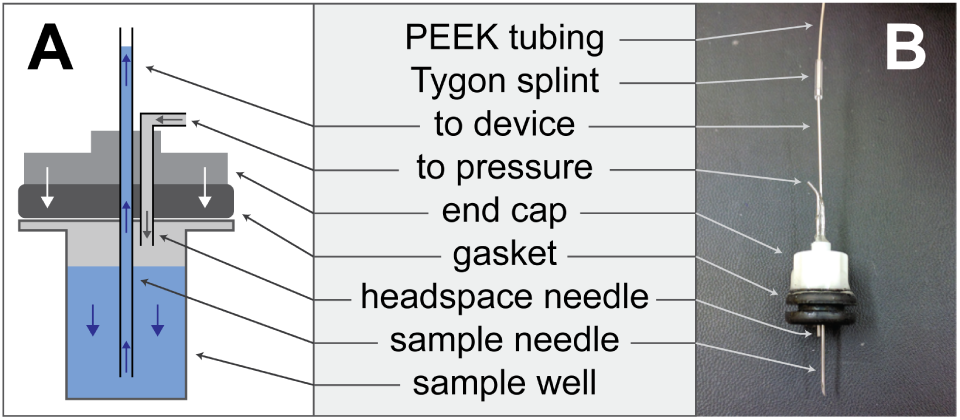
Diagram **(A)** and corresponding image **(B)** showing layout of the dual-lumen ‘sipper’ needle. A rubber gasket seals the needle assembly against a multiwell plate, allowing air pressure applied to the headspace to drive fluid into the sample needle and subsequently onto the device.

#### 3.2.1 Dual-lumen needle with sealing gasket

In conventional positive-pressure microfluidics, sample fluids are carried from pressurized, airtight sample reservoirs to devices via tubing that connects the reservoir sample volume and the device inlet. Adapting this scheme to plate-based wells requires a way to temporarily seal and pressurize each well of interest. We designed a dual-lumen needle that includes one short needle to push air into the well’s headspace and a second longer needle to sample the pressurized fluid volume (Fig. 3). To seal the well, we incorporated a gasket into the needle assembly such that the clamping force of the Z-assembly is sufficient to seal the well and allow pressurization (Fig. 3). An adhesive foil plate seal, commonly used to protect samples and prevent evaporation during techniques such as polymerase chain reaction (PCR), also aids in forming an airtight connection with the needle gasket, while an acrylic brace plate (Fig. 2A: k) prevents the 96-well plate from being deformed by the clamping force of the Z-assembly.

#### 3.2.2 Horizontal needle positioning

To allow iterative sampling, multiwell plates are mounted on an XY-assembly (Fig. 2A: j,k) that translates the plate beneath the dual-lumen needle. The implementation pictured here uses an ASI MS-2000 XY-stage salvaged from a decommissioned Illumina GAIIx sequencer (eBay, $1500) for low-cost construction. However, the AutoSipper can be implemented with any stage that provides sufficient travel and resolution to allow repeatable addressing of all required well positions during an experiment (*e.g.* many standard microscope stages). As the travel of many XY-stages is limited to approximately the dimensions of standard multiwell plates, the attachment points of the XY- and Z-assemblies to the structural frame can be adjusted so that all positions on the stage deck are within the limits of travel.

To ensure that samples remain stably fixed in place, multi-well plates are attached to the XY-stage via 3D-printed deck components (files compatible with the ASI MS-2000 are provided in the micrIO GitHub repository;^†^ see Fig. 2A: k). The primary deck components form a slot which accommodates any ANSI/SLAS standard multiwell plate. An optional 4 × 20-mL rack holds scintillation vials that can help reduce cross-contamination between injections by holding solvents to wash the exterior of the dual-lumen needle and by collecting waste backflushed through the input tubing.

#### 3.2.3 Vertical needle positioning

Once positioned above the desired well, the AutoSipper must compress and seal the well with the dual-lumen needle to sample the well’s contents before raising the needle again and moving to the next location. To facilitate this, the 3D-printed arm bearing the dual-lumen needle is mounted on a carriage that moves along a linear way affixed to a laser-cut acrylic backplate bolted to the 80/20 frame (Fig. 2A: e,f,c,b). The needle arm is moved up and down this linear way by a stepper motor with an integrated controller-driver that drives a lead-screw with ∼100 mm of travel (Lin Engineering CO-4118S-09) (Fig. 2A: e,c,a). The 3D-printed arm bearing the dual-lumen needle (Fig. 2A: e) uses minimal material to fabricate while retaining the structural rigidity necessary to compress the gasket against a well without deflection (design files available in the micrIO GitHub repository^†^). The acrylic backplate has multiple mounting slots to enable flexible vertical and horizontal placement of the Z-assembly along the 80/20 vertical rail (Fig. 2A: b).

#### 3.2.4 Fluidic sampling and washing

To drive sample flow, the headspace needle must be connected to a pressure source capable of pressurizing the well and inducing an application-appropriate flow rate (in our case, up to 12 psi). This can be a constant pressure source, as fluid will only be pushed through tubing when the dual-lumen needle gasket creates a seal with the multiwell plate surface. For applications that are particularly sensitive to cross-contamination (*e.g.* sampling of DNA for downstream PCR applications), scintillation vials for holding wash solvents and collecting backflush waste can be placed on the XY-stage deck, enabling vigorous cleansing of device lines and fluidic components between injections.

### 3.3 Fraction Collector

Multiplexed sample output leverages the full capabilities of on-chip automation to prepare separate fractions and collect each in a separate output vial. Similar to the AutoSipper, the Fraction Collector positions the outlet line directly above the selected well by using an XY-stage to translate the destination plate beneath the outlet tubing (Fig. 2B). However, the Fraction Collector lacks the need for a motorized Z-assembly because it relies on gravity to deposit effluent into destination wells. To flush the outside surface of the outlet tubing and minimize ‘hanging drops’ that could produce cross-contamination, the Fraction Collector features a simple sheath assembly in which the effluent outlet is coaxially centered within a 3D-printed sheath nozzle that allows a low-pressure stream of fluid to rinse the edges of the tubing (*e.g.* with an ethanol-water wash) (Fig. 2B: g,h,i). This sheath flow can be left on at all times or can be temporally activated using available pneumatics control hardware (as in our implementation).

### 3.4 Software

Both the AutoSipper and Fraction Collector may be programmed by our open-source Python package for hardware control, known as *acqpack*, which also enables programming of devices such as microfluidic control manifolds, MFCS units, and syringe pumps that must be coordinated with during an experiment. Up-to-date versions of *acqpack* are installable both via GitHub and pip, aiding in easy deployment and portability.

When building *acqpack*, we noted that users expect a high-level instrument such as an autosampler or fraction collector to have a particular set of properties and functions regardless of the precise hardware implementation. We therefore designed our associated software with a modular architecture that allows users to build an autosampler or fraction collector with alternate hardware so long as they wrap manufacturer-specific hardware commands within a low-level class to provide a standard interface for higher-level classes (Fig. 4). Each device that interfaces with the controlling PC (*i.e.* the Z-motor and the XY-stage controller) has a corresponding low-level class that exposes the commonly used functions of that device. These classes map hardware-level (often serial) commands provided by the hardware manufacturer onto functions that standardize the notion of what a particular object can do (*e.g.* a stage can ‘home’ itself, ‘move’ by a relative number of units, or ‘move’ to an absolute position). High-level classes coordinate low-level functions and provide additional functionality not intrinsic to the lower-level classes. For instance, a high-level *Autosampler* subsumes both a low-level *AsiController* (XY) and *Motor* (Z) while also managing emergent functionality expected of an *Autosampler* (such as coordinate frames and platemaps).

**Fig. 4.**
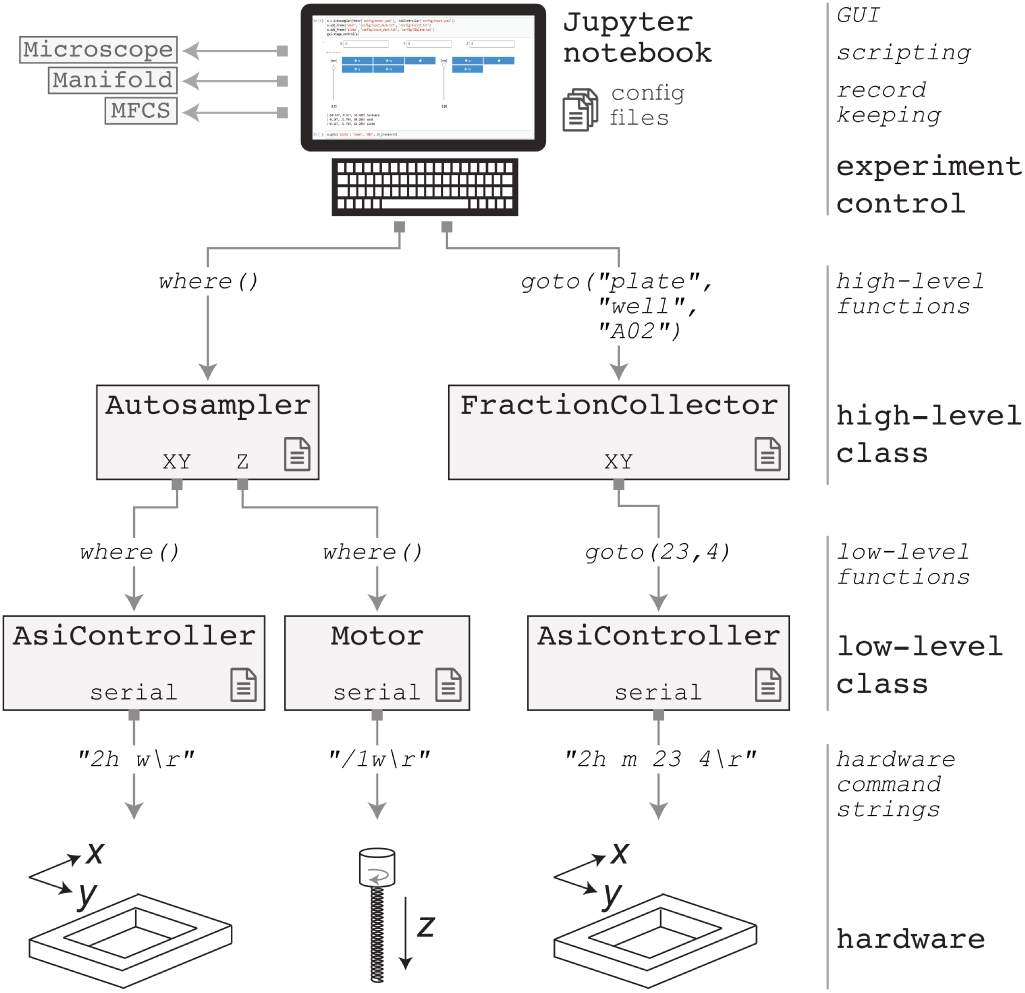
Software architecture of the *acqpack* package. The evolution of two high-level commands (issued from a GUI or script) are shown as they propagate down to hardware.

**Fig. 5.**
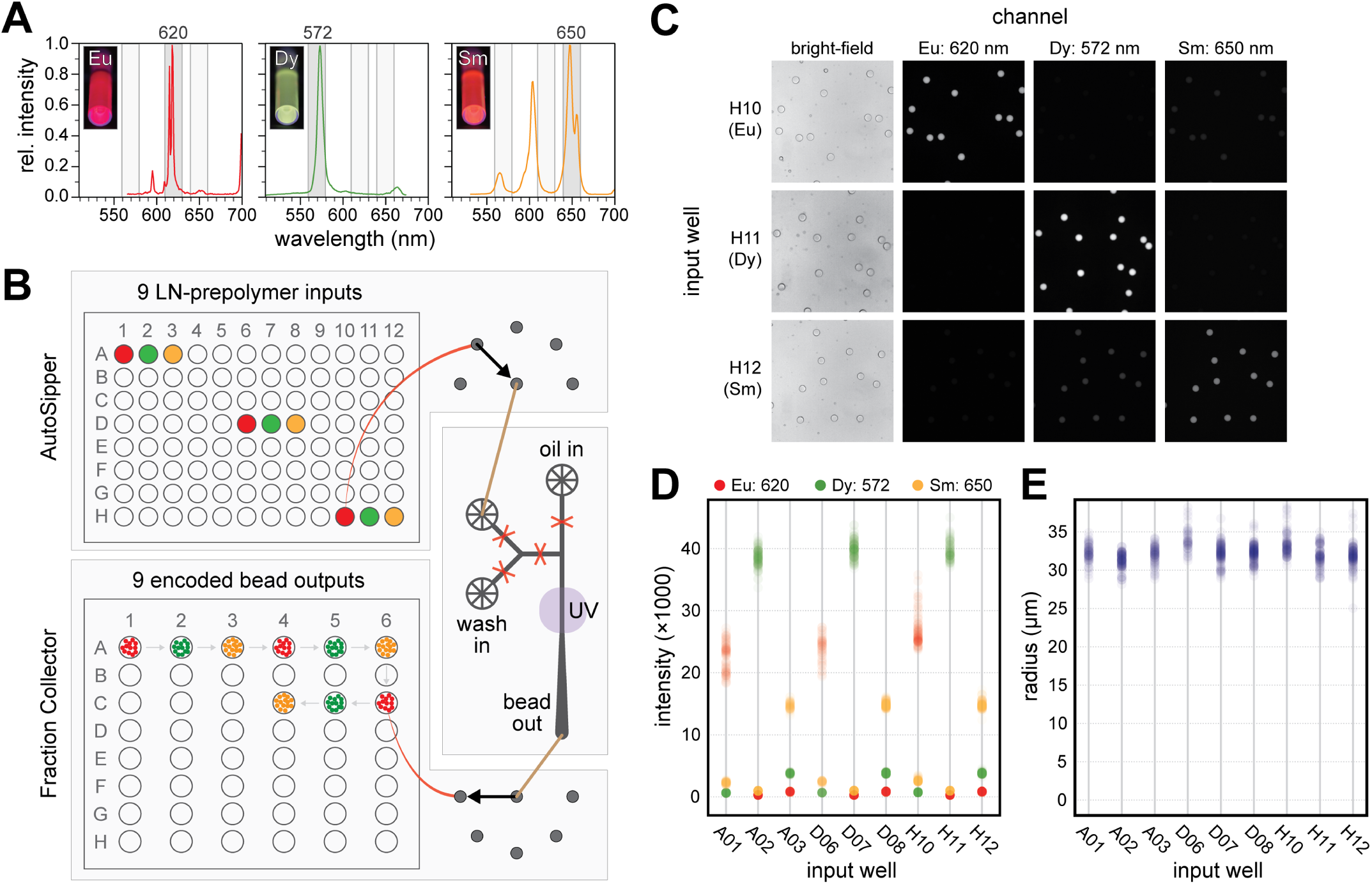
Validation experiment of the AutoSipper and Fraction Collector using spectrally encoded bead synthesis as the test application. **(A)** Photographs (inset) and emission spectra for 3 lanthanide yttrium orthovanadate nanophosphor (LN) species under UV excitation (292 nm): Europium (left), Dysprosium (middle), and Samarium (right). The bandpass filter regions used to distinguish these LNs (620 nm, 572 nm, and 650 nm, respectively) are shown in the background. **(B)** Cartoon showing experimental setup. LN-prepolymer mixtures positioned in 9 wells across a 96-well plate were introduced into a microfluidic droplet generator via the AutoSipper. The droplets produced on-device were polymerized into solid beads via exposure to UV light. Each batch of beads was output to individual wells of a multiwell plate using the Fraction Collector. **(C)** Example multi-channel images of beads from wells H10, H11, and H12 containing putative Eu, Dy, and Sm beads, respectively. **(D)** Per-channel intensity distributions for each well (894 beads total). **(E)** Distribution of bead sizes for each well.

Properly addressing vessels placed on the deck of an autosampler or fraction collector requires reconciling the hardware’s notion of coordinates with those of the vessels. To aid in this point registration and calibration process, *acqpack* allows users to specify the number of rows and columns of vessels (*e.g.* wells) within an array (*e.g.* plate) placed on the deck, along with the hardware coordinates of 3-4 vessels in the array so that the orientation and spacing of the array can be determined. The software then calculates the hardware coordinates of all vessels in the array and saves a platemap: a flat table of vessels, their array indices, and and their hardware coordinates. The user may extend this platemap table with additional columns defining vessel properties, such as contents and names, that then become a valid means to address vessels (*e.g.* a *FractionCollector* can be instructed to *goto* a ‘waste’ vessel). The software also allows users to define and save alternate coordinate frames as transformation matrices (*e.g.* so that a plate may have its own origin, scaling, and orientation).

In addition to coordinate frame and platemap files, parameters for a particular device, such as the serial communication port and settings, are stored in a .yaml configuration file that is loaded when a class is instantiated (rather than being hard-coded into source). This separates the procedural code from the configuration of a particular setup, enabling portability from system to system. To aid users in configuration and calibration of a new AutoSipper or Fraction Collector, associated software documentation and tutorial Jupyter notebooks guide users through setup. Graphical user interfaces (GUIs) for the AutoSipper and Fraction Collector further assist in this manual process.

While Python is all that is required for scripted control of an *Autosampler* and/or *FractionCollector*, Jupyter notebooks provide huge benefits to researchers in all stages of the experimental process that spans protocol development, execution, and analysis. ^26,27^ In the development stage, Jupyter’s cell-based format enables researchers to rapidly test and modify snippets of code and assemble them into a full protocol notebooks. These notebooks can be documented with inline, formatted text cells that allow other researchers to understand and modify the code as necessary. During protocol execution, the notebook gives a detailed record of the code used to execute the experiment which a researcher can further annotate with inline specifics and observations. Finally, the researcher can seamlessly transition to analysis of data collected during execution simply by adding additional cells to process the data and generate inline plots. Similar to traditional notebooks, Jupyter notebooks are useful tools for researchers to communicate what they did and what they discovered.

### 3.5 Platform validation: synthesis of spectrally encoded beads

Demonstrating the utility of the AutoSipper and Fraction Collector requires showing that the platform can reliably: (a) sample from all wells of a 96-well input plate, (b) form a stable pressure seal to push samples through to a microfluidic device, (c) collect samples into specified output vials located throughout a collection rack, (d) perform these tasks without cross-contamination between samples, and (e) do all of these tasks in a programmable, automated manner.

Microfluidic bead generation provides an excellent test application for the AutoSipper and Fraction Collector. First, many labs across the world generate droplets using microfluidic T-junction or flow focusing devices for a variety of applications, ^28^ including single-cell genomics, ^29^ high-throughput protein screening, ^30,31^ and digital droplet PCR. ^32,33^ Second, generation of microfluidic droplets containing spectrally distinct materials allows detection of minute amounts of carryover during a) input, which would manifest as a drift in bead code, and b) output, which would manifest as beads of a certain code being collected into the wrong vial. Lastly, monodisperse droplet formation requires stable input pressure and flow, and so bead size distribution can serve as a proxy for the ability of the AutoSipper to form a robust connection with the sample well. ^34^ We therefore fabricated a microfluidic droplet generator in which an aqueous flow meets an oil flow at a simple T-junction, causing the aqueous flow to pinch off and form droplets. ^22^ The aqueous flow channel includes 2 inlets controlled by on-chip pneumatic valves, making it possible to prime the device with a particular material by directing flow from the inlet to a waste outlet just before the T-junction (Fig. 5B, waste outlet not depicted); similarly, this architecture allows for stringent channel washing between injections by directing flow from the wash inlet to the waste outlet.

To generate spectrally encoded beads, we employed 3 species of LNs (Eu:YVO_4_, Sm:YVO_4_, and Dy:YVO_4_) that can be excited at a single wavelength (292 nm) yet emit visible light in well-separated spectral bands, making it easy to distinguish one from another by imaging with specific bandpass filters (Fig. 5A). To demonstrate that the AutoSipper can reliably sample across its mechanical range, each LN-prepolymer mixture was deposited in triplicate within wells spaced across a 96-well input plate. We then programmed the AutoSipper to sample from each of these wells in series and push the LN-prepolymer mixture through the microfluidic device to form prepolymer droplets. These droplets were polymerized on-chip with a UV light source, and the Fraction Collector deposited the resulting bead batches into separate collection vials in serpentine fashion (Fig. 5B).

To reduce carryover, the AutoSipper was programmed to cleanse the sample line and needle between injections by directing a wash solvent to flow from the bead generator wash inlet back through the AutoSipper inlet tubing. The outside of the dual-lumen needle was also cleaned by dipping into separate scintillation vials containing strong (isopropyl alcohol) and weak (water) wash solvent, and the outlet line was given ample time to clear itself of remaining beads. Once collected, we imaged the beads deposited in each well under bright-field illumination and under UV excitation with emission filtered via 3 bandpass filters to discriminate between the 3 LN species (Fig. 5C). Image analysis of ∼100 beads per each well (894 total) revealed that all beads contained the correct LN without any outlier droplets, establishing a lower limit of cross-contamination of ∼0.1% (Fig. 5D). The beads also demonstrated a reasonably tight size distribution, with an overall coefficient of variation of ∼4.2%, as well as little well-to-well variability (Fig. 5E).

## 4 Conclusions

The potential throughput of microfluidics is often limited by the technical challenges associated with multiplexing and demultiplexing off-chip inputs and outputs, thereby constraining the number of samples and conditions that can be probed. In this paper, we present an open-source hardware and software platform that addresses this challenge by providing direct compatibility between standard multiwell plates and simple microfluidic devices. This setup is easy to build, relatively low-cost, and easily configurable. In this implementation, we build on recent efforts to leverage decommissioned Illumina sequencers for low-cost automation and hardware sourcing; ^35–42^ however, the modular software architecture makes it possible to substitute any mechatronic component so long as its hardware-level commands have been wrapped in a low-level Python class to provide a consistent interface. While several open-source autosamplers and fraction collectors have been developed for applications such as spectroscopy and gas chromatography, ^43–45^ the field of microfluidics has relatively few examples of autosampler and fraction collector usage, with the majority of these examples employing commercial solutions. ^46–50^ More recent work has demonstrated a fully automated, low-cost open-source autosampler that uses gravity-driven flow to feed microfluidic devices, ^51^ as well as a gasketed dual-lumen injector for pressure-driven microfluidic sampling that was similar to our solution but with a manual XY-stage. ^33,52–56^ The micrIO platform merges these two advances into an open-source platform for fully automated, pressure-driven multiwell plate sampling while additionally providing fraction collection capabilities and Python software to coordinate with other devices commonly used in microfluidics. To encourage widespread adoption, we provide extensive documentation and build information (see the micrIO GitHub repository^†^).

The micrIO platform is compatible with a wide variety of simple PDMS devices, including valveless designs, and it requires only a mechanism to drive fluid flow. Here, we have integrated the AutoSipper and Fraction Collector with a microfluidic device designed for pressure-driven flow via a regulated pressure source (*e.g.* a computer-controlled regulator, a voltage-controlled solenoid valve, or regulated house air source). This pressure-driven flow configuration shares infrastructure with devices containing integrated on-chip pneumatic valves, allowing fully automated long-term operation with complete flushing of sample lines and device channels between sample loading. However, we anticipate the AutoSipper and Fraction Collector will also be broadly useful for simple valveless devices that use syringe pumps to set fluid flow rates, especially in light of several recently published open-source syringe pump builds. ^34,57–60^ In this configuration, the dual-lumen needle is vented to ambient pressure (or replaced with a conventional needle) and the device itself is mounted on the AutoSipper deck. After moving the AutoSipper to the correct well location, the syringe pump is used to withdraw fluid from well and into the tubing connected to the needle. After fluid loading, the AutoSipper moves to insert the needle into a device inlet and the syringe pump is switched to drive the loaded fluid from the tubing into the well. This capability is enabled by the high precision and repeatability of the XY-stage and has the potential to greatly simplify droplet generation screens and workflows for high-throughput single-cell sequencing applications, among others. ^61^ By adding a simple microscope and computer vision to a similar syringe pump-driven setup, the platform could further function as an automated colony or cell picker. ^62^ The flexibility of micrIO to meet different challenges in lab is enabled by the modular, open-source nature of its control software and build, and we hope that the community will continue to expand its utility through the design of additional deck components and adaptation of the software to control alternate hardware.

## 5 Author contributions

S.A.L and P.M.F. conceptualized the platform and validation experiment and contributed equally to the preparation of the manuscript. S.A.L. designed and built the platform, coded the hardware control software, performed the validation experiment, and coded the image analysis pipeline. P.M.F. provided funding, resources, mentorship, and project supervision.

## 7 Acknowledgements

This work was supported by NIH grants 1DP2GM123641 and R01GM107132. P.M.F. is a Chan Zuckerberg Biohub Investigator and acknowledges the support of a Sloan Research Foundation Fellowship. The authors would like to thank C. Layton for helpful discussions concerning Illumina GAIIx components, K. Brower for guidance with spectrally encoded bead synthesis, and A. K. White for guidance with bead synthesis, bead imaging, and manuscript revision.

## Footnotes

† Electronic Supplementary Information (ESI) available: [GitHub repository: https://github.com/FordyceLab/micrio]. See DOI: 10.1039/b000000x/

